# Endosomal explosion induced by hypertonicity

**DOI:** 10.1101/730648

**Authors:** Shupeng Wang, Song Wu

**Author notes:** Corresponding author, (S. Wu).

## Abstract

Transferring DNA into cells to regulate cell function is a novel research field in recent decades.
Chitosan is a gene vector with the properties of low-cost and safe, but high efficient delivery has remained challenging. We developed a strategy termed EEIH for endosomal explosion induced by hypertonicity, in which short-time exposure to hypertonic solutions triggers endosomes destabilization like explosions. EEIH can force chitosan/DNA polyplexes to break through the endosomal barriers to approach the nucleus, which results in boosting the transfection efficiency of chitosan in several cell lines. We demonstrate that EEIH is a significant and practical strategy in chitosan transfection system without sophisticated modification of chitosan.

## 1. Introduction

The introduction of specific genes into the cell nucleus has raised considerable interest in quite a long time, both in research and the clinic. Non-viral gene vector transfection is relatively safe compared with viral delivery. Chitosan is safe and degradable as a non-viral gene vector, but the transfection efficiency is limited[1, 2]. Modification of chitosan with arginine[3], histidine[4], lysine[4], Polyethylenimine [5], polyethylene glycol (PEG)[1], lipid[6], inorganic nanoparticles[7], and branched structure[8, 9] can improve chitosan transfection to a certain degree, but the resulting materials are still not efficient enough or mass produced, which limits the wide range of applications.

PEGylated gene carriers generally achieve higher gene transfection efficiency[1, 10, 11]. PEG could promote transfection by another method. In 1985, Gopal demonstrated that at the end of incubating cells with DEAE-dextran and DNA, the treatment with a 40% PEG3400 solution for 2-4 min could improve transfection efficiency effectively. Based on this finding, we examined whether brief exposure to PEG200, PEG1000 and PEG6000 could improve transfection efficiency at the end of treatment of cells with chitosan/DNA polyplexes. We found the treatment of PEG of lower molecular weight enhanced transfection more effectively, which suggested high osmotic pressure may play a nonnegligible role.

In this study, we describe a strategy to boost chitosan transfection. We demonstrate that the briefly hypertonic treatment of cells incubated with chitosan/DNA polyplexes offers superior gene transfection efficiency. We term this strategy “EEIH” for endosomal explosion induced by hypertonicity. The mechanism may be that the hypertonicity treatment can change the intracellular osmotic pressure, and then endosomes destabilize like explosions, and foreign DNA escapes from endosomes, which lead to foreign DNA enter into the nucleus to boost transfection (Fig. 1). Our results highlight two points, the first is that EEIH is easy to operate, chitosan can achieve satisfied transfection efficiency combination with EEIH for used widely, the second is that EEIH may force exogenous substance uptaken to escape from endosomes to disperse in the cytosol, which has great promise in gene transfection and drug delivery system.

**Fig. 1.**
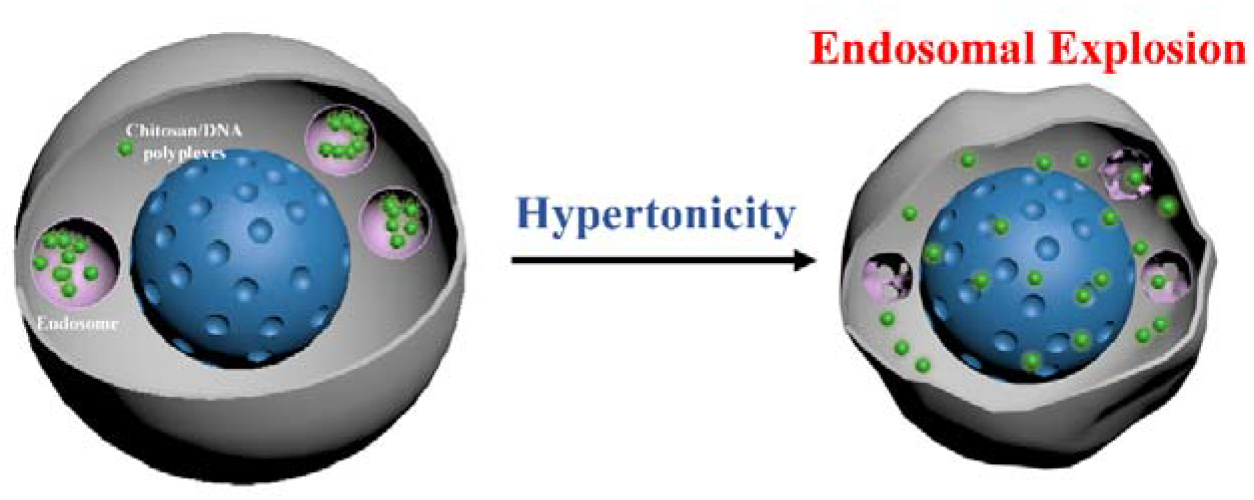
Schematic illustration showing the endosomal explosion induced by hypertonicity

## 2. Materials and Methods

### 2.1 Materials and cell lines

Chitosan was purchased from Aladdin (China). Glycerol, dimethylsulfoxide (DMSO), PEG200, PEG1000 and PEG6000 were purchased by Sigma (USA). YOYO-1, LysoTracker Red and Hoechst 33342 were purchased by Invitrogen (USA). Sodium chloride (NaCl) was purchased from Sangon Biotech (China). The chemicals were used as received without further purification. Plasmid DNA pCMV-C-EGFP(5010 bp, Cat. No. D2626) was purchased by Beyotime Biotechnology (China).

HEK293T (a human embryonic kidney cell line, ATCC) and HeLa (a human cervical cancer cell line, ATCC) were cultured in Dulbecco’s Modified Eagle Medium (DMEM) (Gibco, USA) supplemented with 10% fetal bovine serum and 1% penicillin-streptomycin. 5637 (a bladder cancer cell line, ATCC) was cultured in RPMI-1640 medium (Gibco, USA) supplemented with 10% fetal bovine serum and 1% penicillin-streptomycin.

### 2.2 Preparation of chitosan/DNA polyplexes

A chitosan solution (200 μg/ml, pH 5.5) was prepared as previously described[1]. An equal volume of the DNA solution (100 μg/ml in molecular biology grade water) and the sterile-filtered (0.22 mm) chitosan solution were mixed and vortexed for 30 s. The chitosan/DNA polyplexes were incubated at room temperature for 20 min before further use.

### 2.3 In vitro transfection

The transfection media for chitosan (DMEM, 5 mM MES, pH 6.5) was prepared for further use, according to previous studies[12, 13]. The cells were seeded at 7.5×10^4^ cells/well in 24-well plates for 24 h before gene transfection, followed by the medium in each well was replaced with 500 μl of the transfection media, Chitosan/DNA polyplexes were added into each well and incubated with the cells at the dose of 0.5 μg DNA/well. After 4 h at 37 □ and 5% CO_2_, the transfection media was replaced with 250ul 1x PBS containing different concentrations (weight/volume percentage) of compounds, followed by exposed treatment at room temperature for 3 min. Then the cells were washed twice with PBS and then replaced with complete media for another 42 h before transfection efficiency analysis. The expression of EGFP in cells was observed by fluorescent microscopy (Olympus, Japan) and analyzed by flow cytometry (BD, USA).

## 3. Results and discussion

### 3.1. The effect of PEG treatment on chitosan transfection

In an early study, Gopal found briefly exposed to a 40% solution of PEG3400 had increased the transfection efficiency of DEAE-dextran, but the mechanism has not illuminated clearly[14]. To validate the effect of the high concentration of PEG on chitosan transfection, we used different molecular weight and concentration of PEG to treat the Hela cells incubated with chitosan/DNA polyplexes. 3-min PEG treatment had a distinct influence on chitosan transfection (Fig. 2). It is worth noting that lower molecular weight PEG achieved higher transfection efficiency at the same concentration, so we speculated that osmolality might influence transfection efficiency.

**Fig. 2.**
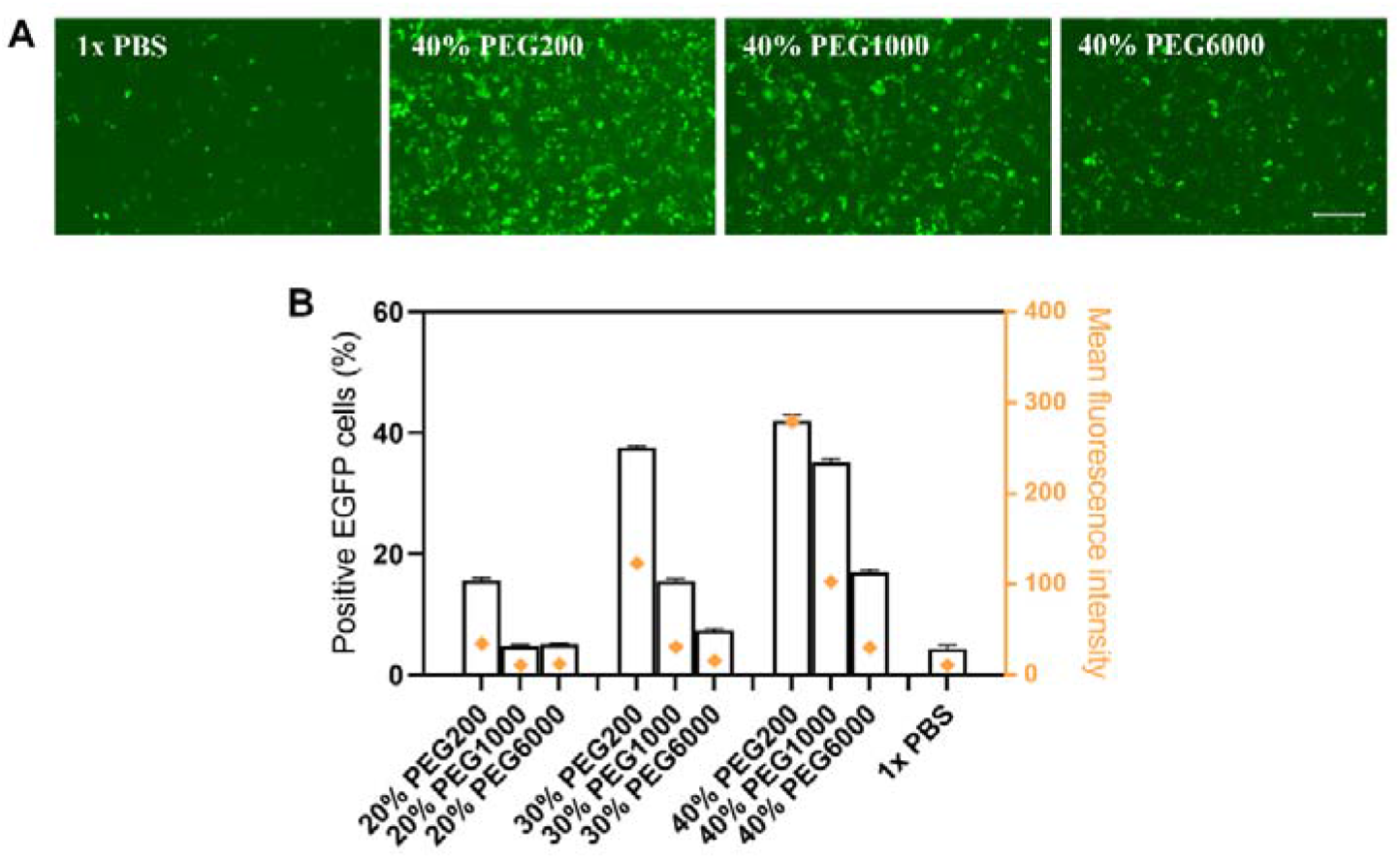
The effect of PEG treatment on chitosan transfection in Hela cells. (A) Fluorescence microscopy images of Hela cells transfected (scale bar, 500 μm). (B) Positive EGFP cells (%) and mean fluorescence intensity were recorded by flow cytometry. Data are indicated as mean ± SD (n = 3).

### 3.2. The effect of glycerol or DMSO treatment on chitosan transfection

To further examine the role of osmolality, we explored the impact of the hypertonic solution induced by glycerol or DMSO on chitosan transfection. As shown in Fig. 3, the results of high transfection efficacy were confirmed in HeLa cells, HEK293T cells and 5637 cells.

**Fig. 3.**
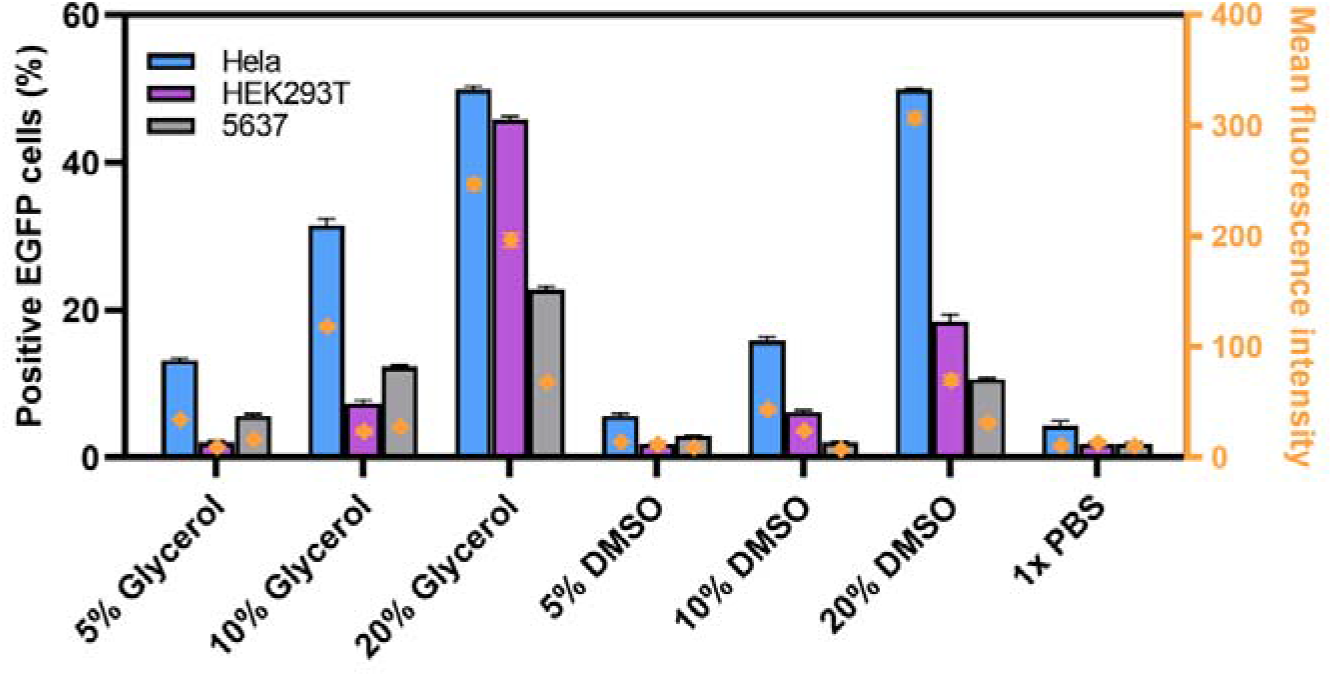
The effect of different concentration of glycerol or DMSO solutions on the chitosan in Hela, HEK293T and 5637 cells. Positive EGFP cells (%) and mean fluorescence intensity were recorded by flow cytometry. Data are indicated as mean ± SD (n = 3).

Based on demonstrated above, we noticed that glycerol, DMSO and low molecular weight PEG were permeable cryoprotectants[15]. We wondered whether the high transfection efficiency was mainly due to the process of permeable compounds diffused into cells.

### 3.3 The effect of NaCl hypertonicity treatment on chitosan transfection

Next, we tested whether hypertonicity induced by the compound without the ability to penetrate cell membranes could improve chitosan/DNA polyplexes transfection efficiency. We examined the effect of NaCl hypertonicity treatment on chitosan transfection. As shown in Fig. 4, 3-min exposed to 10% NaCl solution improves transfection efficacy significantly. Here, we speculated that hypertonicity played the main role in high chitosan transfection rather than the effect of permeable compounds.

**Fig. 4.**
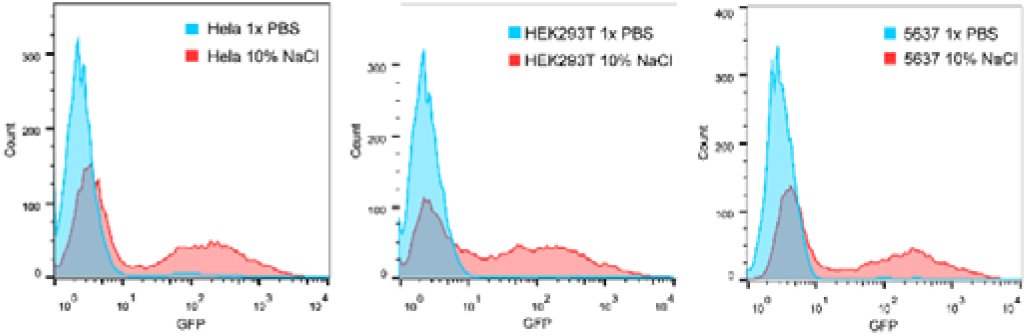
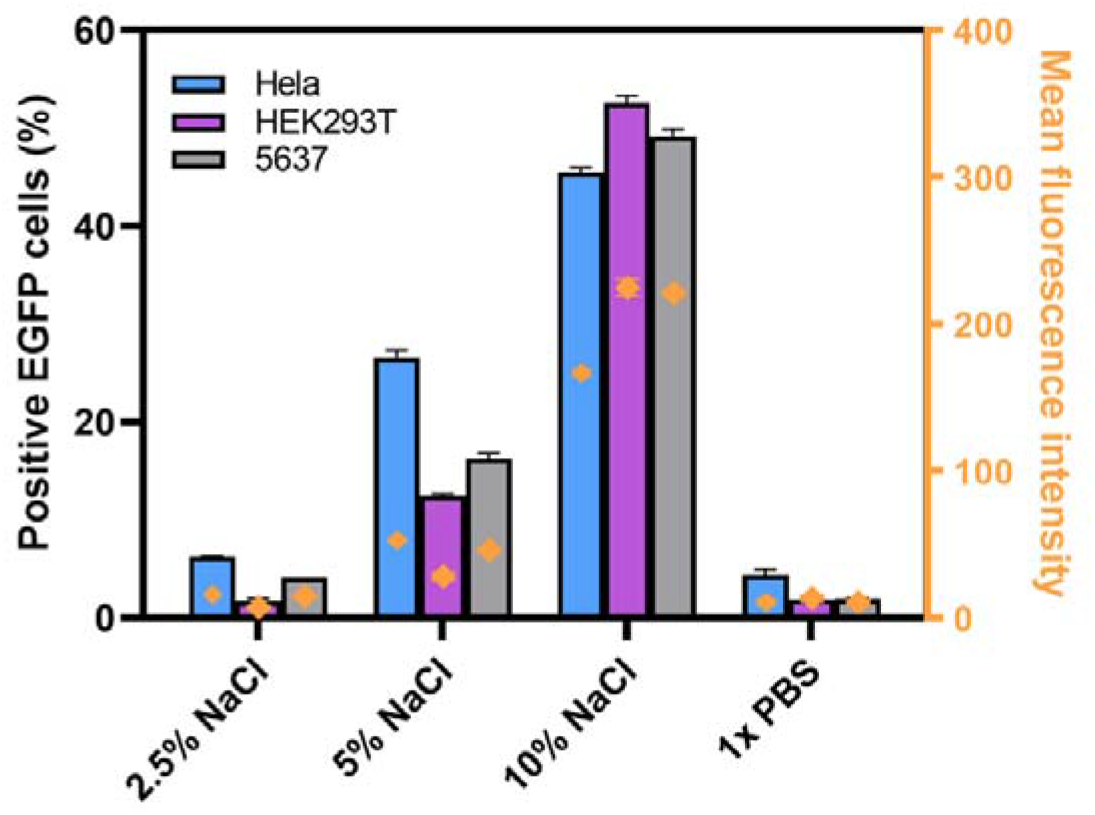
The effect of NaCl hypertonicity treatment on chitosan transfection in Hela, HEK293T, and 5637 cells. Positive EGFP cells (%) and mean fluorescence intensity were recorded by flow cytometry. Data are indicated as mean ± SD (n = 3).

## 4. Conclusion

Here we present a low-cost, highly efficient and flexibly adaptable method of chitosan transfection. Endosomal escape is a key step in the transfection process. In the hypertonic condition, the EEIH strategy can destabilize endosomes, facilitate chitosan/DNA polyplexes endocytosed to escape from the endosomes into the nucleus. This strategy can be used as a versatile tool in the development of gene and drug delivery, especially for the therapy of hollow organs such as the cervix uteri and the bladder.

